# Branched actin polymerization drives invasive protrusion formation to promote myoblast fusion during skeletal muscle regeneration

**DOI:** 10.1101/2024.09.30.615960

**Authors:** Yue Lu, Tezin Walji, Pratima Pandey, Chuanli Zhou, Christa W. Habela, Scott B. Snapper, Rong Li, Elizabeth H. Chen

**Affiliations:** Department of Molecular Biology, University of Texas Southwestern Medical Center, Dallas, TX, USA; Department of Immunology, University of Texas Southwestern Medical Center, Dallas, TX, USA; Department of Neurology, Johns Hopkins University School of Medicine, Baltimore, MD, USA; Department of Pediatrics, Boston Children’s Hospital, Boston, MA, USA; Department of Cell Biology, Johns Hopkins University School of Medicine, Baltimore, MD, USA; Mechanobiology Institute, National University of Singapore, Singapore, Singapore; Department of Cell Biology, University of Texas Southwestern Medical Center, Dallas, TX, USA; Hamon Center for Regenerative Science and Medicine, University of Texas Southwestern Medical Center, Dallas, TX, USA; Harold C. Simmons Comprehensive Cancer Center, University of Texas Southwestern Medical Center, Dallas, TX, USA

## Abstract

Skeletal muscle regeneration is a multistep process involving the activation, proliferation, differentiation, and fusion of muscle stem cells, known as satellite cells. The fusion of <u>s</u>atellite <u>c</u>ell-derived mononucleated <u>m</u>uscle cells (SCMs) is indispensable for the generation of multinucleated, contractile myofibers during muscle repair. However, the molecular and cellular mechanisms underlying SCM fusion during muscle regeneration remain poorly understood. In this study, we uncovered an essential role for branched actin polymerization in SCM fusion. Using conditional knockouts of the Arp2/3 complex and its actin nucleation-promoting factors, N-WASP and WAVE, we demonstrated that branched actin polymerization is required for the SCM fusion, but not for satellite cell proliferation, differentiation, and migration. We showed that the N-WASP and WAVE complexes have partially redundant functions in regulating SCM fusion. Furthermore, we revealed that branched actin polymerization is essential for generating invasive protrusions at the fusogenic synapses in SCMs. Taken together, our study has identified new components of the myoblast fusion machinery in skeletal muscle regeneration and demonstrated a critical role for branched actin-propelled invasive protrusions in this process.

## Introduction

Skeletal muscle is a unique tissue composed of elongated multinucleated cells known as myofibers (Frontera and Ochala 2015). In response to injury, skeletal muscle has the capacity to repair injured myofibers in a process called muscle regeneration. Muscle regeneration is dependent on the resident muscle stem cells, known as satellite cells (Yin et al. 2013; Relaix et al. 2021). Satellite cells are located between the myofiber plasma membrane and the basement membrane (BM), the latter of which is a layer of extracellular matrix material composed of collagen, glycoproteins, and proteoglycans (Seale et al. 2000; Webster et al. 2016). Satellite cells express high levels of Pax7, which is a paired domain-containing transcription factor, and remain quiescent under normal conditions (Seale et al. 2000; Le Grand and Rudnicki 2007; Cheung and Rando 2013; Yin et al. 2013). Upon injury, satellite cells are activated and then proliferate and differentiate into fusion-competent muscle cells to repair the injury (Le Grand and Rudnicki 2007; Cheung and Rando 2013; Yin et al. 2013; Hindi and Millay 2022). Once the satellite cell-derived mononucleated muscle cells, which will be referred to as SCMs hereafter, fill the space within the BM remnants, known as ghost fibers, they would fuse with each other and/or with injured myofibers to regenerate the muscle (Webster et al. 2016; Collins et al. 2024). SCM fusion occurs rapidly between day 3.5 to 5 post injury (dpi) and persists till ∼dpi 10 (Collins et al. 2024). Despite the importance of SCM fusion in skeletal muscle regeneration, the molecular and cellular mechanisms underlying SCM fusion during muscle regeneration remain poorly understood. To date, only two proteins, the bi-partite myoblast fusogens myomaker (MymK) (Millay et al. 2013) and myomixer (MymX)/myomerger/minion (Bi et al. 2017; Quinn et al. 2017; Shi et al. 2017; Zhang et al. 2017), have been shown to be required for SCM fusion *in vivo* (Millay et al. 2014; Bi et al. 2018). Identifying additional components of the SCM fusion machinery will not only facilitate our understanding of muscle regeneration but also provide more options to enhance muscle regeneration upon injury.

Studies in multiple organisms have provided significant insights into the evolutionarily conserved mechanisms underlying myoblast fusion during embryogenesis (Chen 2011; Kim et al. 2015a; Schejter 2016; Deng et al. 2017; Kim and Chen 2019; Lee and Chen 2019; Petrany and Millay 2019). It has been demonstrated that embryonic myoblast fusion in *Drosophila*, zebrafish and mouse embryos is mediated by an invasive podosome-like structure composed of actin-propelled membrane protrusions at the fusogenic synapse (Sens et al. 2010; Luo et al. 2022; Lu et al. 2024). The branched actin nucleator, the Arp2/3 complex (Richardson et al. 2007; Berger et al. 2008), and its actin nucleation-promoting factors (NPFs), the Neural Wiskott Aldrich Syndrome Protein (N-WASP) (Massarwa et al. 2007; Schäfer et al. 2007; Sens et al. 2010; Gruenbaum-Cohen et al. 2012), and WASP-family verprolin-homologous protein (WAVE) (Schroter et al. 2004; Richardson et al. 2007; Gildor et al. 2009; Sens et al. 2010), are required for generating the invasive protrusions at the fusogenic synapse. Additional actin cytoskeletal regulators upstream of the NPFs also function in mammalian myoblast fusion during development, such as activators for N-WASP (Cdc42) and WAVE (Rac1) (Vasyutina et al. 2009), and the bi-partite guanine nucleotide exchange factor for Rac1 (Dock180 [also known as Dock1] and Elmo) (Laurin et al. 2008; Tran et al. 2022). A subunit of the WAVE complex (Nap1) has been shown to promote myoblast fusion in cultured C2C12 myoblasts (Nowak et al. 2009). Previous studies have shown actin-propelled protrusions between cultured SCMs (Randrianarison-Huetz et al. 2018), as well as membrane protrusions at the fusion sites of cultured SCMs (Eigler et al. 2021). Recent studies have revealed the mechanism underlying the formation of invasive protrusions at the fusogenic synapses – it takes the coordination of two Arp2/3 NPFs (WAVE and N-WASP) and two actin-bundling proteins (dynamin and WASP-interacting protein (WIP)) to generate mechanically stiff actin bundles that propel invasive protrusions (Zhang et al. 2020b; Lu et al. 2024). The essential function of the actin cytoskeleton in myoblast fusion has been further highlighted by the fact that each of the bi-partite muscle fusogens, MymK and MymX, requires a functional actin cytoskeleton to induce myoblast fusion (Millay et al. 2013; Zhang et al. 2017).

Despite all the previous studies, the potential function of branched actin polymerization in muscle regeneration *in vivo* has not been directly tested. Here, using satellite cell-specific knockout (KO) mice of Arp2/3 and NPFs, we show that branched actin polymerization is indispensable for muscle regeneration. In particular, Arp2/3 and NPFs are required for the formation of invasive protrusions during SCM fusion, but not satellite cell proliferation, differentiation or migration. Thus, we have identified new components of the SCM fusion machinery *in vivo* and demonstrated a critical role for branched actin-propelled invasive protrusions in skeletal muscle regeneration.

## Results

### SCMs populate the ghost fibers after macrophage departure at early stages of skeletal muscle regeneration

To examine SCMs after injury, we injured the tibialis anterior (TA) muscles by BaCl_2_ injection (Fig. 1A), and labeled the differentiating SCMs using an antibody against NCAM, a cell adhesion molecule highly expressed in these cells (Capkovic et al. 2008), and ghost fibers using an antibody against Laminin, a major component of the BM (Webster et al. 2016) (Fig. 1B). Since macrophages are present in the ghost fibers to clear the necrotic debris of the damaged myofibers (Collins et al. 2024), we also labeled macrophages with an antibody against MAC-2, a member of the lectin family expressed on the cell surface of macrophages (Hohsfield et al. 2022). At dpi 2.5, the differentiating SCMs and macrophages were two major cell populations residing within the ghost fibers, occupying 39.9% and 47.8% of the total volume, respectively. By dpi 3.5, SCMs filled 98.2% of the ghost fiber volume, whereas macrophages only accounted for 1.8% (Fig. 1C), with most of the macrophages residing in the interstitial space outside of the ghost fibers (Fig. 1B), which would account for the high overall number of macrophages in the regenerating muscle tissues in this time period (Collins et al. 2024). Of note, our confocal (Fig. S1A, S1B, and Supplementary Video 1 & 2) and transmission electron microscopy (TEM) (Fig. S1C) analyses of regenerating TA muscles at dpi 3 also revealed narrow openings (∼1 μm diameter) on the BM of the ghost fibers, through which macrophages (MAC-2^+^) with a ∼20 μm diameter (as a round cell) were traversing, suggesting that macrophages enter and/or escape the ghost fibers by squeezing through tiny openings on the BM. By dpi 4.5, most of the SCMs have fused into multinucleated primary myofibers (Fig. 1B) (Collins et al. 2024).

**Figure 1.**
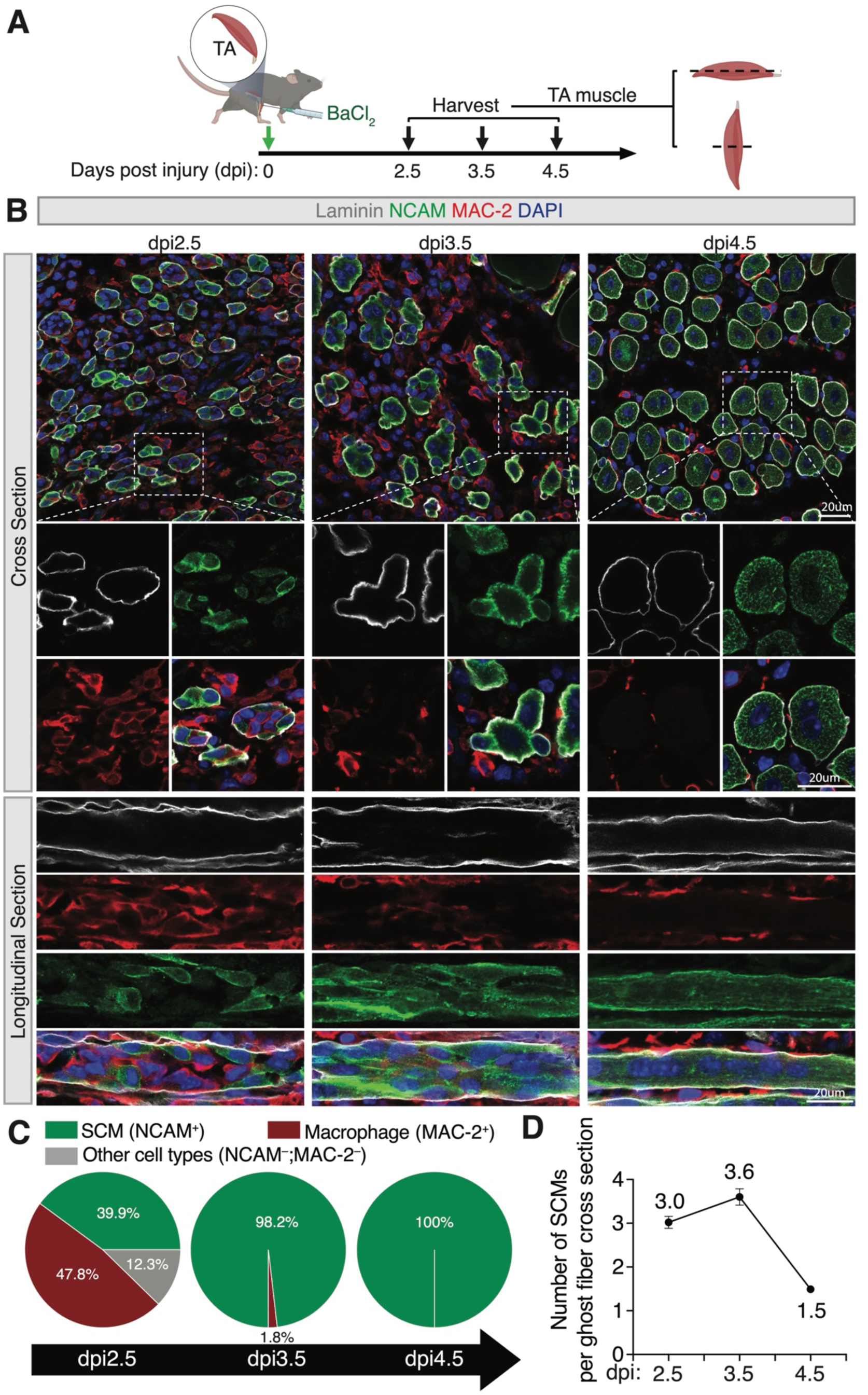
Spatiotemporal coordination of macrophages and SCMs during skeletal muscle regeneration. **(A)** Diagram of the TA muscle injury scheme. The TA muscles of the wild-type mice were injured by intramuscular injection of BaCI_2_. The injured TA muscles were collected at dpi 2.5, 3.5, and 4.5 for cross and longitudinal sectioning and immunostaining. **(B)** Immunostaining with anti-Laminin, anti-NACM, and anti-MAC-2 of the cross and longitudinal sections of TA muscles at the indicated time points. Note the decrease in the macrophage number within the ghost fiber at dpi 3.5 (compared to dpi 2.5), and the fusion of SCMs between dpi 3.5 and 4.5. Scale bars: 20 μm. **(C)** Quantification of the percentage of macrophages and differentiated SCMs within ghost fibers at the indicated time points. *n* = 3 mice were analyzed for each time point and > 98 ghost fibers in each mouse were examined. Mean ± s.d. values are shown. **(D)** Quantification of the number of differentiated SCMs in a single cross section of a ghost fiber at indicated time points. *n* = 3 mice were analyzed for each time point and > 98 ghost fibers in each mouse were examined. Mean ± s.e.m values are shown.

### Branched actin polymerization is required for mammalian skeletal muscle regeneration

Given that the Arp2/3 complex-mediated branched actin polymerization is required for myoblast fusion during mouse embryogenesis (Lu et al. 2024), we asked whether the Arp2/3 complex is required for skeletal muscle regeneration in adults. Toward this end, we generated satellite cell-specific, tamoxifen-inducible KO mice for ArpC2, a subunit of the Arp2/3 complex (Goley and Welch 2006), by breeding *Pax7*^CreERT2^ mice (Lepper et al. 2009) with *ArpC2*^fl/fl^ mice (Wang et al. 2016). The conditional knockout (cKO) mouse line *Pax7*^CreERT2^; *ArpC2*^fl/fl^ will be referred to as *ArpC2*^cKO^ hereafter. The littermates of the *Pax7*^CreERT2^ mice without the floxed *ArpC2* allele were used as wild-type controls. To induce genetic deletion of *ArpC2* in satellite cells, we performed intraperitoneal injection of tamoxifen to the control and mutant mice every two days over a period of ten days (Fig. 2A). The *ArpC2* KO in satellite cells was confirmed by western blot (Fig. S2). Satellite cell-specific *ArpC2* KO did not affect TA muscle weight and size in uninjured muscle (Fig. S3A-S3C). However, muscle injury by BaCl_2_ resulted in a significant decrease (87.7 **±** 2.0%) in the cross-sectional area (CSA) of regenerated myofibers in *ArpC2*^cKO^ mice compared to their littermate controls at dpi 14 (Fig. 2B and 2C) and dpi 28 (Fig. S4A and S4B). Consistent with this, the frequency distribution of CSA displayed a significant shift toward the small size in the mutant mice (Fig. 2D). Taken together, these data demonstrate that the Arp2/3-mediated branched actin polymerization is essential for skeletal muscle regeneration.

**Figure 2.**
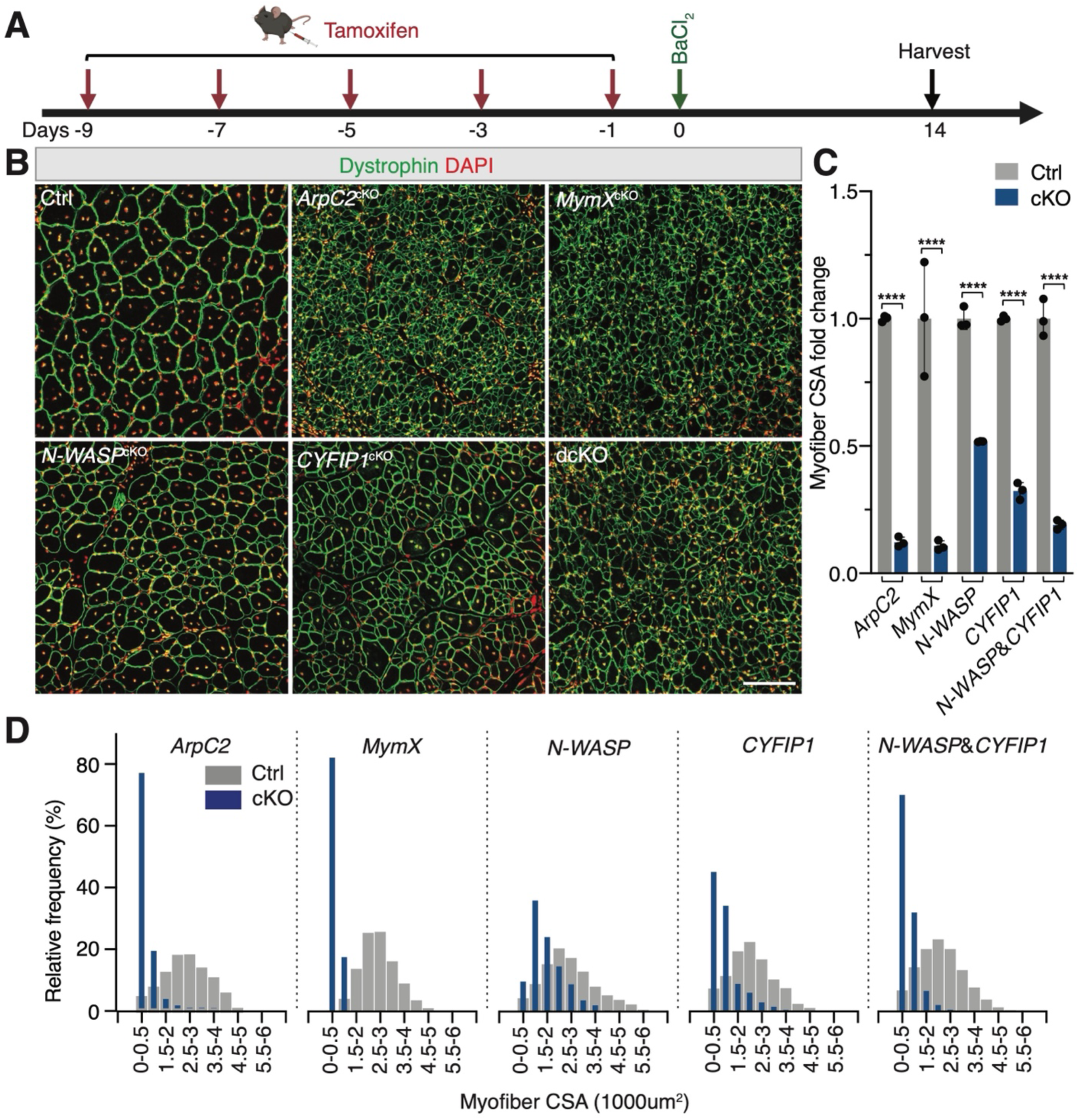
Branched actin polymerization is required for skeletal muscle regeneration. (A) Schematic diagram of tamoxifen and BaCI_2_ treatment and subsequent CSA analysis at dpi 14. (B) Dystrophin and DAPI staining of the cross sections of TA muscles at dpi 14 from the control (Ctrl) and mutant mice. Note that the myofiber CSA is moderately decreased in *N-WASP*^cKO^ and *CYFIP1*^cKO^ mice, and severely reduced in dcKO, *ArpC2*^cKO^ and *MymX*^cKO^ mice. Scale bar: 100 μm. (C) The fold change of myofiber CSA in mutant mice *vs*. control mice. *n* = 3 mice were analyzed for each time point and > 200 fibers in each mouse were examined. Mean ± s.d. values are shown in the bar graph, and significance was determined by two-tailed student’s t-test. ****: p < 0.0001. (D) Frequency distribution of myofiber CSA of TA muscles in the control and mutant mice at dpi 14. *n* = 3 mice of each genotype were examined and > 200 ghost fibers in each mouse were examined.

### Branched actin polymerization is required for SCM fusion

To pinpoint the specific step of skeletal muscle regeneration – satellite cell proliferation, differentiation, migration, and SCM fusion – in which branched actin polymerization is required, we performed immunostaining using antibodies that specifically mark these steps. As shown in Fig. S5A-S5E, the percentages of muscle cells positive for the proliferation marker (Ki67) and the muscle differentiation marker (MyoG) in the injured TA muscles were similar between control and *ArpC2*^cKO^ mice (Fig. S5A-S5E). In addition, live imaging analysis showed that cultured *ArpC2* KO SCMs exhibited normal migration and cell-cell contact behaviors (Supplementary Video 3). These results demonstrate that branched actin polymerization is dispensable for satellite cell proliferation, differentiation, and migration during skeletal muscle regeneration. Thus, the reduced muscle size in the *ArpC2* mutant mice is likely due to defects in SCM fusion.

To monitor the SCM fusion phenotypes, we examined the regenerating TA muscles of the control and *ArpC2*^cKO^ mice at dpi 4.5, when myoblast fusion leading to primary myofiber formation is mostly completed (Figure 1B) (Collins et al. 2024). While most of the SCMs within the ghost fibers had fused in the control animals, the ghost fibers in the *ArpC2*^cKO^ mice contained differentiated (NCAM^+^), but mostly unfused, SCMs, which were readily observed in cross sections, comparable to those in the *MymX*^cKO^ mice (Fig. 3A-3C and Fig. S6A). Consistent with this, the frequency distribution of SCM numbers in a cross section per ghost fiber in the *ArpC2*^cKO^ and *MymX*^cKO^ mice displayed a dramatic shift toward higher numbers compared with that of the wild-type mice (Fig. 3D). These results indicate that the actin cytoskeleton plays as an essential role in SCM fusion as the fusogenic proteins. Interestingly, expression levels of the fusogenic proteins, MymK and MymX, in the TA muscle of *ArpC2*^cKO^ mutant mice were either similar or higher compared to that of wild-type mice (Fig. S5F-S5H), suggesting that the fusion defect in the *ArpC2*^cKO^ mutant mice was not due to a lack of fusogen expression. Consistent with this, cultured *ArpC2* KO SCMs exhibited a severe fusion defect despite undergoing normal differentiation (Fig. 3E-3G) and pharmacologically inhibiting Arp2/3 with CK666 also led to a similar fusion defect (Fig. S6B-S6D). In addition, cell-mixing experiments using wild-type and *ArpC2* KO SCMs showed that *ArpC2* KO SCMs failed to fuse with wild-type cells (Fig. 3H and 3I), indicating that branched actin polymerization is required in both fusing partners. Taken together, these results demonstrate that branched actin polymerization is required in SCMs for their fusion during skeletal muscle regeneration. The severe myoblast fusion defects observed in early stages of regeneration (e.g. dpi 4.5) provide a good explanation for the presence of thin muscle fibers in ArpC2 cKO mice at dpi 14 (Fig. 2B and 2C) and dpi 28 (Fig. S4A and S4B). These thin muscle fibers could be either elongated mononucleated muscle cells or multinucleated myofibers each containing a small number of nuclei due to occasional fusion events (comparable to those in Myomixer cKO muscles) (Fig. 2B and 2C; Fig. S4A and S4B). Whether Arp2/3 and branched actin polymerization may play a role in the growth and/or maintenance of post-fusion multinucleated myofibers requires future loss-of-function studies to inactivate ArpC2 using a myofiber-specific Cre driver.

**Figure 3.**
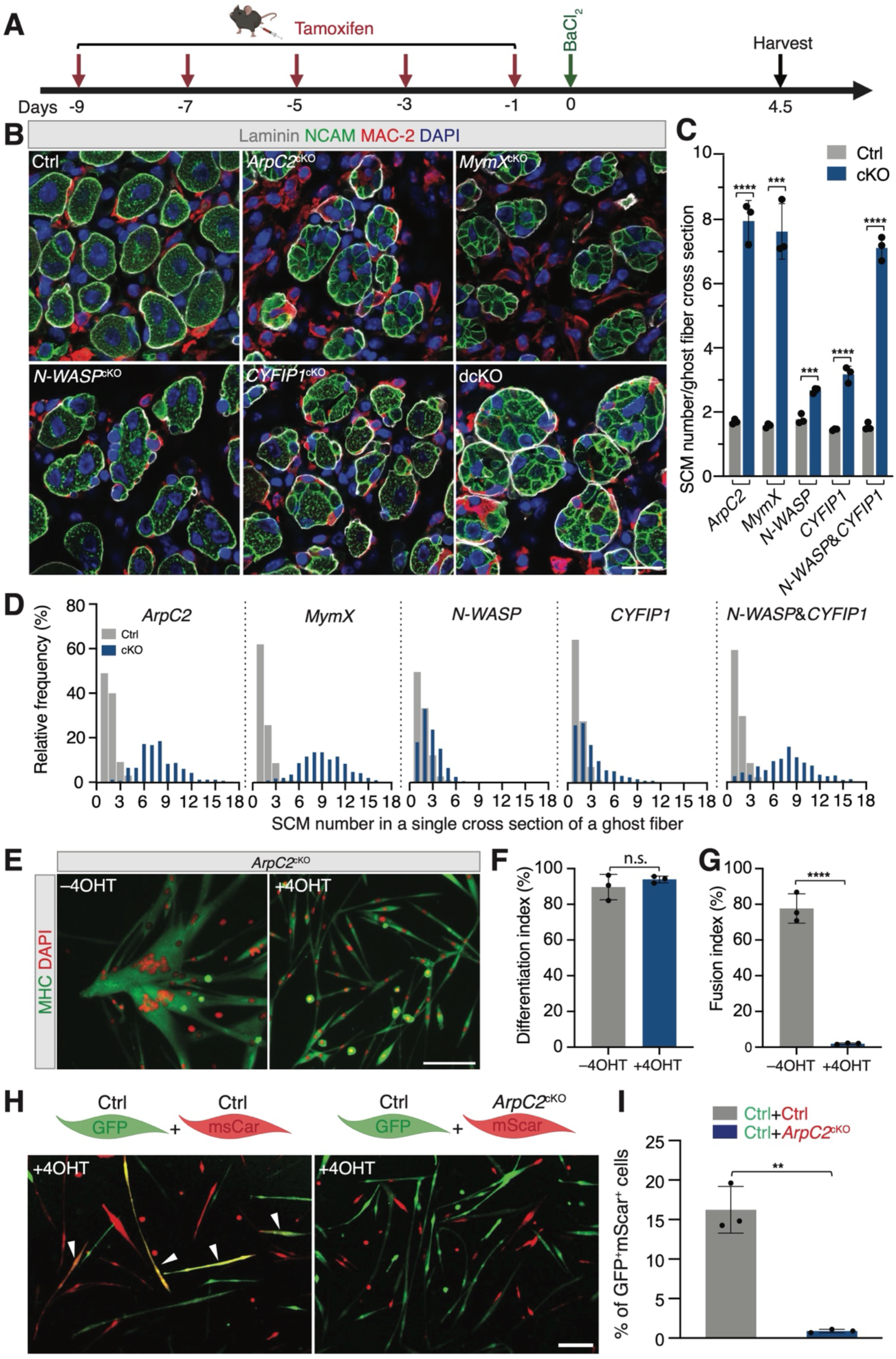
Branched actin polymerization is required for SCM fusion. (A) Schematic diagram of tamoxifen and BaCI_2_ treatment and subsequent SCM number analysis at dpi 4.5. (B) Immunostaining with anti-laminin, anti-NCAM, and anti-MAC-2 of the cross sections of TA muscles at dpi 4.5 from the control and mutant mice. Note that each ghost fiber in the control mice contained 1-2 centrally nucleated myofiber at dpi 4.5, indicating the near completion of SCM fusion. The ghost fibers in *N-WASP*^cKO^ and *CYFIP1*^cKO^ mice contained more SCMs, indicating impaired SCM fusion. Note that even more SCMs were seen in dcKO, *ArpC2*^cKO^ and *MymX*^cKO^ mice. Scale bar: 20 μm. (C) Quantification of the SCM number in a single cross section of a ghost fiber from TA muscles of the control and mutant mice at dpi 4.5. *n* = 3 mice were analyzed for each time point and > 80 ghost fibers in each mouse were examined. Mean ± s.d. values are shown in the bar graph, and significance was determined by two-tailed student’s t-test. ***: p < 0.001 and ****: p < 0.0001. (D) Frequency distribution of SCM number in a single cross section of a ghost fiber from TA muscles of the mutant mice and their littermate control. *n* = 3 mice of each genotype were analyzed and > 80 ghost fibers in each mouse were examined. (E) ArpC2 is required for SCM fusion in cultured cells. The satellite cells isolated from *ArpC2*^cKO^ mice were maintained in GM without or with 2 μM 4OH-tamoxifen (4OHT) for 10 days. Subsequently, the cells were plated at 70% confluence in GM. After 24 hours, the cells were cultured in DM for 48 hours, followed by immunostaining with anti-MHC and DAPI. Note the robust fusion of the control (–4OHT) SCMs and the severe fusion defects in *ArpC2* KO (+4OHT) SCMs. Scale bar: 100 μm. **(F**, **G)** Quantification of the differentiation index (% of nuclei in MHC^+^ cells *vs*. total nuclei) and fusion index (% of nuclei in MHC^+^ myotubes with ≥ 3 nuclei *vs*. total nuclei) of the two types of cells shown in (E). *n* = 3 independent experiments were performed. Mean ± s.d. values are shown in the bar graphs, and significance was determined by two-tailed student’s t-test. ****: p < 0.0001; n.s: not significant. (**H**) ArpC2 is required in both fusion partners. Fluorescence images from cell-mixing experiments using differentially labelled SCMs are shown. The satellite cells isolated from *ArpC2*^cKO^ mice were infected with retroviruses encoding GFP or mScarleti (mScar). Next, the GFP^+^ cells were maintained in GM for 10 days (Ctrl GFP^+^ cells), and the mScar^+^ cells were maintained in GM without (Ctrl mScar^+^ cells) or with 2 μM 4OH-tamoxifen (*ArpC2*^cKO^ mScar^+^ cells) for 10 days. Subsequently, the Ctrl GFP^+^ cells were mixed with Ctrl mScar^+^ cells or with *ArpC2*^cKO^ mScar^+^ cells with a ratio of 1:1 and plated at 70% confluence in GM. After 24 hours, the cells were cultured in DM for 48 hours followed by direct fluorescent imaging. Arrowheads indicate syncytia derived from both GFP and mScar cells. Scale bar: 100 μm. (**I**) Percentage of GFP^+^mScar^+^ syncytia in total cells shown in (H). Mean ± s.d. values are shown in the bar graph, and significance was determined by two-tailed student’s t-test. **: p < 0.01.

### N-WASP and WAVE families have partially redundant functions in regulating SCM fusion

Activation of the Arp2/3 complex requires the actin NPFs, including the WASP and WAVE family of proteins (Goley and Welch 2006). To examine their potential functions in mammalian muscle regeneration, we generated single and double cKO mice for N-WASP [the WASP family member with high expression in SCMs (Lu et al. 2024)] and CYFIP1 [a subunit of the WAVE complex (Eden et al. 2002)], respectively. The cKO mouse line *Pax7*^CreERT2^*; N-WASP*^fl/fl^ will be referred to as *N-WASP*^cKO^, *Pax7*^CreERT2^; *CYFIP1*^fl/fl^ as *CYFIP1*^cKO^, and *Pax7*^CreERT2^; *N-WASP*^fl/fl^; *CYFIP1*^fl/fl^ as dcKO hereafter. Target protein knockouts in SCMs were confirmed by western blot (Fig. S2).

For the single cKO mice, immunostaining revealed a moderate but significant reduction of TA myofiber CSA at dpi 14 by 48.2 ± 0.1% in *N-WASP*^cKO^ and 67.7 ± 3.3% in *CYFIP1*^cKO^ mice, respectively, compared to their littermate controls (Fig. 2B and 2C). The myofiber CSA of dcKO mice further decreased to 80.9 ± 1.8%, comparable to the 87.7 ± 2.0% observed in the *ArpC2*^cKO^ mice (in which both N-WASP and WAVE complexes are defective) and to the 89.3 ± 1.9% in the *Mymx*^cKO^ mice (Fig. 2B and 2C). Moreover, the *N-WASP*^cKO^ and *CYFIP1*^cKO^ single KO mice exhibited moderate myoblast fusion defects at dpi 4.5, which were exacerbated in dcKO mice (Fig. 3B-3D), despite normal satellite cell proliferation and differentiation, as well as the persistent fusogenic protein expression in the dcKO mice (Fig. S5A-S5H). Thus, our data revealed partially redundant functions between N-WASP and WAVE NPFs in promoting myoblast fusion during skeletal muscle regeneration.

### Branched actin polymerization promotes invasive protrusion formation during SCM fusion

To investigate the mechanism by which branched actin polymerization regulates SCM fusion during muscle regeneration, we first examined the cellular structure at the fusogenic synapse of cultured SCMs. Live cell imaging of cultured SCMs expressing Arp2-mNeongreen (mNG) and LifeAct-mScarleti (mScar) at day two in differentiation medium (DM) revealed Arp2- and F-actin-enriched finger-like protrusions projecting from the invading cells into their fusion partners (receiving cells) at the fusogenic synapse prior to cell membrane fusion (Fig. 4A and Supplementary Video 4). Consistent with this, TEM analysis of the TA muscle at dpi 3.5 in wild-type mice revealed finger-like protrusions projected by SCMs invading their neighboring cells (20.3 ± 16.5% of SCMs exhibited invasive protrusions, n = 83 SCMs from 20 ghost fibers examined) (Fig. 4B and 4C). The average length and width of the invasive finger-like protrusions were 422 ± 200 nm and 121 ± 73 nm, respectively (Fig. S7A and S7B, n = 32 invasive protrusions examined). In contrast, muscle cells in *ArpC2*^cKO^ mice seldom p invasive protrusions (0.5 ± 2.2% of SCMs exhibited invasive protrusions, n = 147 SCMs from 20 ghost fibers examined) (Fig. 4B), whereas protrusions in SCMs of *MymX*^cKO^ mice appeared normal (Fig. 4B and 4C, 24.1 ± 15.6% of SCMs exhibited invasive protrusions, n = 93 SCMs from 20 ghost fibers examined; Fig. S7A and S7B, n = 29 invasive protrusions examined). Therefore, branched actin polymerization, but not the fusogenic protein MymX, is required for invasive protrusion formation to promote myoblast fusion during adult muscle regeneration.

**Figure 4.**
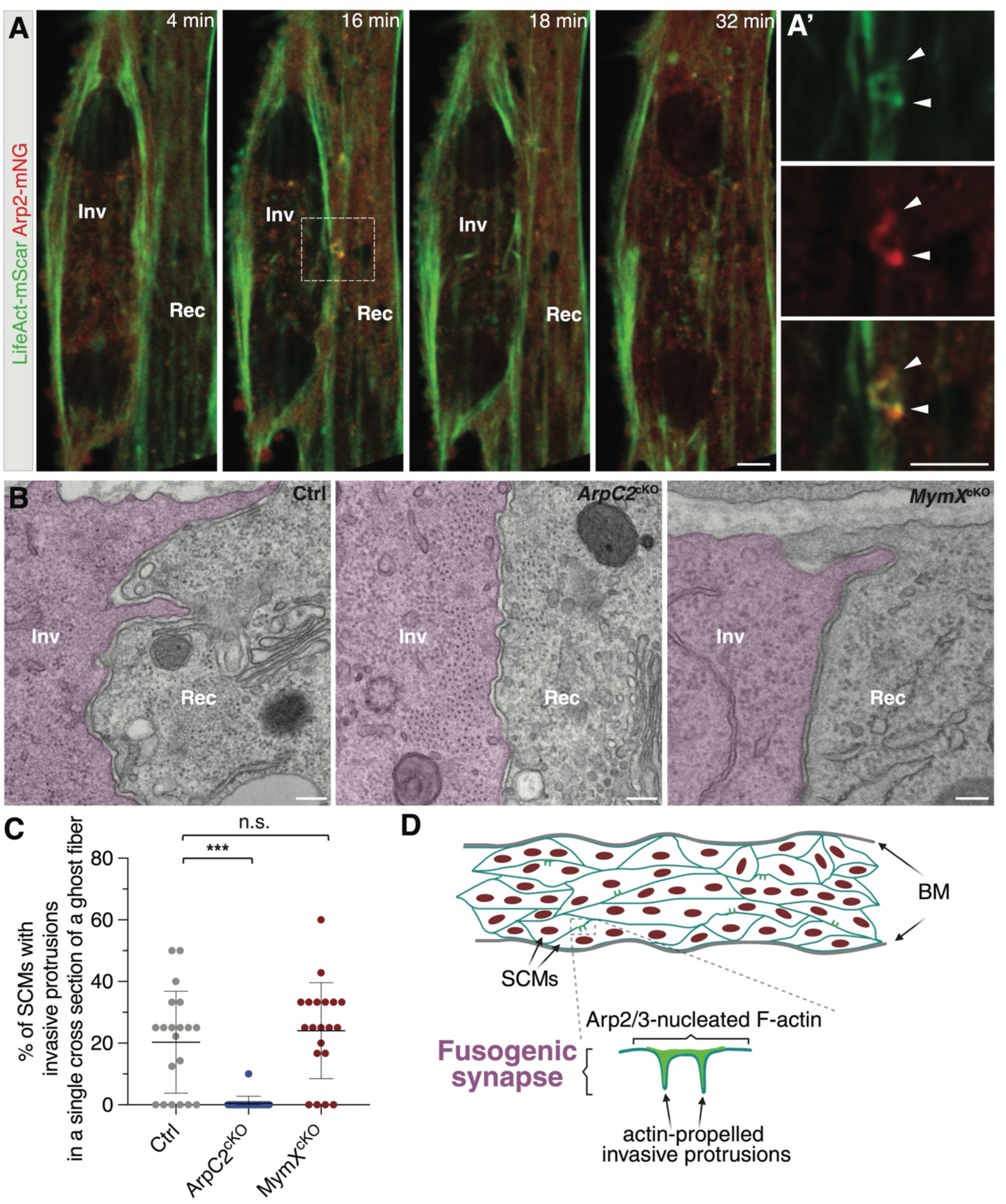
Branched actin polymerization is required for invasive protrusion formation during SCM fusion. **(A)** Still images of a fusion event between two LifeAct-mScar and Arp2-mNG co-expressing SCMs (see Supplemental Video 4). The boxed area is enlarged in (A’). Note the presence of two invasive protrusions (16 minute, arrowheads) enriched with LifeAct-mScar and Arp2-mNG at the fusogenic synapse. *n* = 8 fusion events were observed with similar results. Scale bar: 5 μm. **(B)** TEM of TA muscle cells in wild-type control, *ArpC2*^cKO^, and *MymX*^cKO^ mice at dpi 3.5. The invading SCMs are pseudo-colored in light magenta. Note the finger-like protrusions projected by SCMs invading their neighboring cells in control and *MymX*^cKO^, but not in the *ArpC2*^cKO^, mice. Scale bars: 500 nm. **(C)** Quantification of the percentage of SCMs with invasive protrusions in a single cross section of a ghost fiber in the mice with genotypes shown in (B) at dpi 3.5. At least 83 SCMs from n = 20 ghost fibers in each genotype were quantified. Mean ± s.d. values are shown in the dot plots, and significance was determined by two-tailed student’s t-test. ***p < 0.001; n.s.: not significant. **(D)** A model depicting the function of Arp2/3-mediated branched actin polymerization in promoting invasive protrusion formation to promote SCM fusion during skeletal muscle regeneration. BM: basement membrane.

## Discussion

In this study, we show that the Arp2/3 complex-mediated branched actin polymerization is indispensable for SCM fusion, but not for satellite cell proliferation, migration, or differentiation during muscle regeneration. The Arp2/3 NPFs, N-WASP and WAVE, exhibit partially redundant functions in regulating SCM fusion. Our live cell imaging and electron microcopy analysis revealed actin-propelled invasive protrusions at the fusogenic synapses of SCMs, and our genetic analysis demonstrated a requirement for branched actin polymerization in generating these protrusions. Taken together, we propose that branched actin polymerization promotes mammalian muscle regeneration by facilitating the formation of invasive protrusions at the fusogenic synapse (Fig. 4D).

Studies in multiple organisms, including *Drosophila*, zebrafish, and mouse, have demonstrated that myoblast fusion during embryogenesis is mediated by actin-propelled invasive membrane protrusions (Sens et al. 2010; Jin et al. 2011; Duan et al. 2012; Duan et al. 2018; Luo et al. 2022; Lu et al. 2024). These protrusions enhance the plasma membrane contact areas between the fusion partners and increase the mechanical tension of the fusogenic synapse to promote fusion (Chen 2011; Shilagardi et al. 2013; Kim et al. 2015a; Kim et al. 2015b; Kim and Chen 2019; Lee and Chen 2019). The current study has revealed a similar role for invasive protrusions in promoting myoblast fusion during adult skeletal muscle regeneration, demonstrating that the same cell fusion machinery required during embryogenesis is reused in adult muscle regeneration. It is striking that depleting the branched actin polymerization machinery results in a severe SCM fusion defect similar to depleting the fusogenic protein MymX, highlighting the indispensable role for actin cytoskeletal rearrangements in SCM fusion. Indeed, our previous work with a reconstituted cell-fusion culture system led to the discovery that fusogens and branched actin regulators are two minimal components of the cell-cell fusion machinery, and that actin-propelled invasive protrusions are required to bring the two apposing cell membranes into close proximity for fusogen engagement (Shilagardi et al. 2013). It would be interesting to determine whether invasive protrusions promote the trans-interactions of fusogens at the mammalian fusogenic synapse.

## Materials and Methods

### Mouse Genetics

All animals were maintained under the ethical guidelines of the UT Southwestern Medical Center Animal Care and Use Committee according to NIH guidelines. C57BL/6J (stock: 000664) and *Pax7*^CreERT2^ (stock: 012476) (Lepper et al. 2009) mice were obtained from The Jackson Laboratory. The *ArpC2*^fl/fl^ (Wang et al. 2016), *N-WASP*^fl/fl^ (Cotta-de-Almeida et al. 2007), and *CYFIP1*^fl/fl^ (Habela et al. 2020) mice were previously described. The *MymX*^fl/fl^ line (Bi et al. 2018) was generously provided by Dr. Eric N. Olson. The control and mutant male littermates were used in each cohort of experiments.

### Tamoxifen and BaCl_2_-Induced Muscle Injury

Tamoxifen (Sigma; T5648) was dissolved at 20 mg/ml in corn oil. 100 μl tamoxifen/corn oil solution was administered by intraperitoneal injection to 2-month-old male mice as schematized in Figures. To induce muscle injury, BaCl_2_ (Sigma; 342920) was dissolved in sterile saline to a final concentration of 1.2%, aliquoted, and stored at −20°C. Mice were anesthetized by isoflurane inhalation, the legs were shaved and cleaned with alcohol, and TA muscles were injected with 50 μL of BaCl_2_ with a 28-gauge needle.

### Satellite Cell Isolation and Culture

Satellite cells were isolated from limb skeletal muscles of 2-month-old male mice. Briefly, muscles were minced and digested in 800U/ml of type II collagenase (Worthington; LS004196) in F-10 Ham’s medium (ThermoFisher Scientific; 11550043) containing 10% horse serum for 90 minutes at 37 °C with rocking to dissociate muscle fibers and dissolve connective tissues. The dissociated myofiber fragments were collected by centrifugation and digested in 0.5U/ml dispase (Gibco; 17105041) in F-10 Ham’s medium for 30 minutes at 37 °C with rocking.

Digestion was stopped with F-10 Ham’s medium containing 20% FBS. Cells were then filtered from debris, centrifuged and resuspended in growth medium (GM: F-10 Ham’s medium supplemented with 20% FBS, 4 ng/ ml FGF2, 1% penicillin–streptomycin and 10mM HEPEs). The cell suspension from each animal was pre-plated twice on the regular 100 mm tissue culture-treated dishes for 30 minutes at 37 °C to eliminate fibroblasts. The supernatant containing mostly myoblasts was then transferred into collagen-coated dishes for culture in GM. To validate the KO efficiencies of the target genes, skeletal muscle from six mice of each genotype were pooled for satellite cell isolation. To induce myogenic differentiation, satellite cells were cultured in DM (DMEM supplemented with 2% horse serum, 1% penicillin-streptomycin and 10mM HEPEs).

### Pharmacological Treatments of Satellite Cells

To pharmacologically inhibit branched actin polymerization in SCMs, the Arp2/3 inhibitor CK666 (50 μM) were added into the DM at day 0 of differentiation of wild-type SCMs. After 48 hours, the cells were fixed in 4% paraformaldehyde (PFA) and stained with anti-MHC and DAPI to assess their differentiation and fusion index.

To delete *ArpC2* in SCMs *in vitro*, satellite cells isolated from *ArpC2*^cKO^ mice were cultured in GM supplemented with or without 2 μM 4-hydroxytamoxifen (Sigma; H6278) for 10 days. Subsequently, the cells were trypsinized and plated at 70% confluency in DM. After 48 hours, the cells were fixed in 4% PFA and stained with anti-MHC and DAPI to assess their differentiation and fusion index.

### Retroviral Vector Preparations and Expression

The cytosolic GFP, cytosolic mScarleti, LifeAct-mScarleti and Arp2-mNeongreen constructs were described in the previous study (Lu et al. 2024), and assembled into the retroviral vector pMXs-Puro (Cell Biolabs; RTV-012) using the NEBuilder HiFi DNA Assembly Cloning Kit (NEB; E2621L). To package the retrovirus, two micrograms of retroviral plasmid DNA was transfected into platinum-E cells (Cell Biolabs; RV-101) using the FuGENE HD transfection reagent. Two days after transfection, the virus-containing medium was filtered and concentrated with Retro-X Concentrator (Clontech, PT5063-2) following the manufacturer’s protocol. The concentrated retroviruses were diluted in GM (with a 1:1000 dilution), mixed with polybrene (7 μg/ml), and used to infect cells. One day after infection, cells were washed with PBS and cultured in fresh GM.

### Immunohistochemistry

To co-stain NCAM, MAC-2, and Laminin, the 4% PFA fixed TA muscles were dehydrated in 30% sucrose at 4°C overnight. The specimens were embedded in Tissue-Plus O.C.T. Compound (Fisher Scientific; 23-730-571) and 12-μm cryosections were collected onto Superfrost Plus Microscope Slides (Fisher Scientific;12-550-15). Then, the cryosections were incubated with blocking buffer for 20 minutes at room temperature (RT), followed by overnight incubation with rabbit anti-NCAM (1:200; Millipore; AB5032), rat anti-MAC-2 (1:200; Biolegend; 125401), and rat anti-Laminin-2 (1:500; Sigma; L0663) at 4°C. To stain for dystrophin, the freshly dissected TA muscles were snap frozen in Tissue-Plus O.C.T. Compound and 12-μm cryosections were collected onto Superfrost Plus Microscope Slides. Next, the sections were fixed in 4% PFA for 12 minutes at RT, washed three time with PBS and incubated with blocking buffer for 20 minutes at RT, followed by overnight incubation with rabbit anti-dystrophin (1:200; Abcam; ab15277) at 4°C. To co-stain Pax7, MyoG, Laminin, and Ki67, the freshly dissected TA muscles were snap frozen in Tissue-Plus O.C.T. Compound and 12-μm cryosections were collected onto Superfrost Plus Microscope Slides. Then, the sections were fixed in 2% PFA for 5 minutes at RT, washed three time with PBS and incubated with blocking buffer supplemented with M.O.M blocking reagent (1:25; Vector; MKB-2213-1) for 60 minutes at RT, followed by overnight incubation with mouse anti-Pax7 (1:2; DSHB; Pax7), mouse anti-MyoG (1:2; DSHB; F5D), rat anti-Laminin-2 (1:500; Sigma; L0663), and rat anti-Ki67 (1:500; ThermoFisher Scientific; 14-5698-82) at 4°C. After the incubation with primary antibodies, the sections were extensively washed with PBS and then incubated with alexa fluor-conjugated secondary antibodies for one hour at RT. Subsequently, the sections were washed with PBS and subjected to imaging using a Leica TCS SP8 inverted microscope.

### Western Blot

For western blots, proteins were isolated from the cultured SCMs or TA muscle using ice-cold RIPA buffer (150mM NaCl, 1% NP40, 0.1% SDS and 50mM Tris, PH7.4) containing protease and phosphatase inhibitors (Cell Signaling Technologies; 5872) for 20 minutes. The supernatants were collected by centrifugation at 140,000 x g for 15 minutes. Protein concentrations were determined using the Bradford Protein Assay Kit (Bio-Rad; 5000201). 10-30 μg total protein was loaded for each sample and separated by 10% SDS-PAGE gel and transferred to PVDF membranes (Millipore; GVHP29325). Then, the membranes were blocked for one hour at RT in PBS containing 5% nonfat dry milk and 0.1% Tween-20 (PBSBT) and subsequently were incubated with primary antibodies diluted at 1:1000 in PBSBT overnight at 4°C. The membranes were then washed with PBST and incubated with appropriate HRP-conjugated secondary antibodies diluted in PBSBT for one hour at RT. After extensive washes with PBST, the membranes were developed with the ECL western blotting substrate (ThermoFisher Scientific; 32209). The following primary antibodies were used: sheep anti-ESGP/MymX (1:1000; R&D Systems; AF4580), mouse anti-MymK (Zhang et al. 2020a) (1:1000), and rabbit anti-β-Tubulin (1:1000; Cell Signaling Technologies; 2146).

### Time-lapse Imaging and Analysis

Time-lapse imaging of cells incubated in 5% CO_2_ at 37 °C was performed on a Nikon A1R confocal microscope with a Nikon Biostation CT. The satellite cells were seeded on fibronectin-coated cover glass (MATTEK; P35G-0-14-C) and imaged using a 40× (0.4 NA) objective at indicated time points after switching from GM to DM. The cells were imaged at two- or five-minute interval. After time-lapse imaging, ImageJ (NIH, 64-bit Java 1.8.0_172) was used to project the z-stacks in 2D, using maximum intensity projection and the resulting 2D images were assembled into a time-lapse video.

### Electron Microscopy

To observe the invasive protrusions at the contact sites of SCMs during muscle regeneration *in vivo*, TA muscle at dpi 3.5 were fixed in a solution containing 3% paraformaldehyde, 2% glutaraldehyde, 1% sucrose, 3mM CaCl_2_ in 0.1M sodium cacodylate buffer (pH 7.4) overnight at 4°C. Samples were subsequently washed with 0.1M cacodylate buffer containing 3% sucrose and 3mM CaCl_2_, and post fixed with 1% osmium tetroxide in 0.1M sodium cacodylate buffer for 1.5 hours on ice. The muscle samples were stained with 2% uranyl acetate, dehydrated and embedded in EPON resin as previously described (Zhang and Chen 2008). The embedded samples were then cut into 70-nm thick sections using LEICA ultramicrotome (UC6) and collected on copper slot grids. These sections were post-stained with 2% uranyl acetate and sato’s lead solution and examined using a JEOL 1400 transmission electron microscope.

### Statistics and Reproducibility

Statistical significance was assessed using two-tailed student’s t-test. The *P* values were obtained using GraphPad Prism 8. The numbers of biological replicates for each experiment are indicated in the figure legends.

---

**Supplementary Figure 1.**
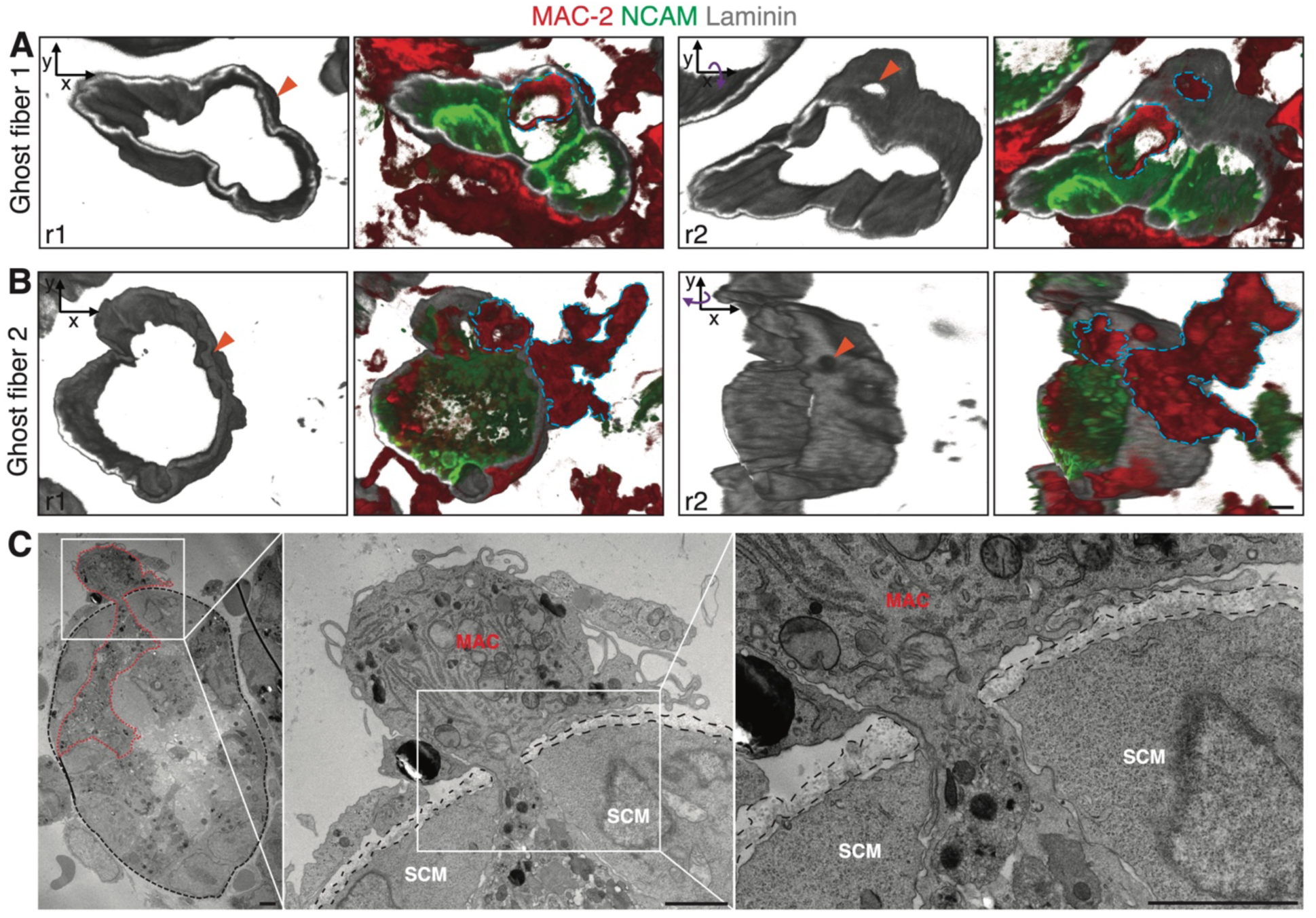
Macrophages extravasate the ghost fibers by traversing the BM. **(A** and **B)** Macrophages traversing the BM of ghost fibers shown by confocal microscopy. TA muscle cross sections of wild-type mice at dpi 3 were immunostained with anti-laminin, anti-NCAM, and anti-MAC-2, followed by confocal imaging. The confocal z-stacks were reconstructed to 3D images. Two examples are shown here. For each traversing macrophage (delineated by cyan dotted lines), images at two rotational angles are shown (r1 and r2). Note the small opening (arrowhead) on the BM through which a macrophage was passing (see supplementary video 1 & 2). **(C)** Macrophages traversing the BM of ghost fibers shown by TEM. The TA muscles as described in (A) and (B) were subjected to TEM processing. The BM is outlined by black dotted lines. The traversing macrophage is delineated by red dotted lines in the left panel. MAC: macrophage; SCM: SC-derived muscle cell. Scale bars: 2 μm.

**Supplementary Figure 2.**
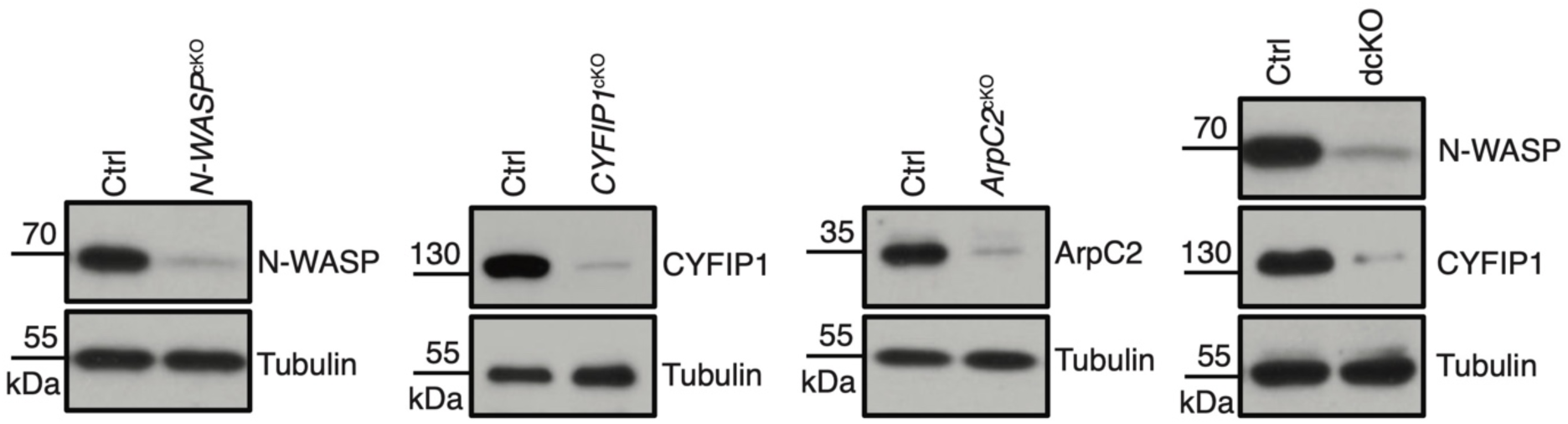
The knockout efficiency of targeted proteins in the mouse models. Mice with indicated genotypes were treated with tamoxifen as described in Figure 1A. The skeletal muscles were isolated and digested one day after the 5^th^ tamoxifen injection. The satellite cells were enriched *in vitro* for two days, yielding a culture composed of > 90% satellite cells, followed by western blot for the targeted proteins. For each genotype, skeletal muscle from *n* = 6 mice were pooled for satellite cell isolation and western blot analysis.

**Supplementary Figure 3.**
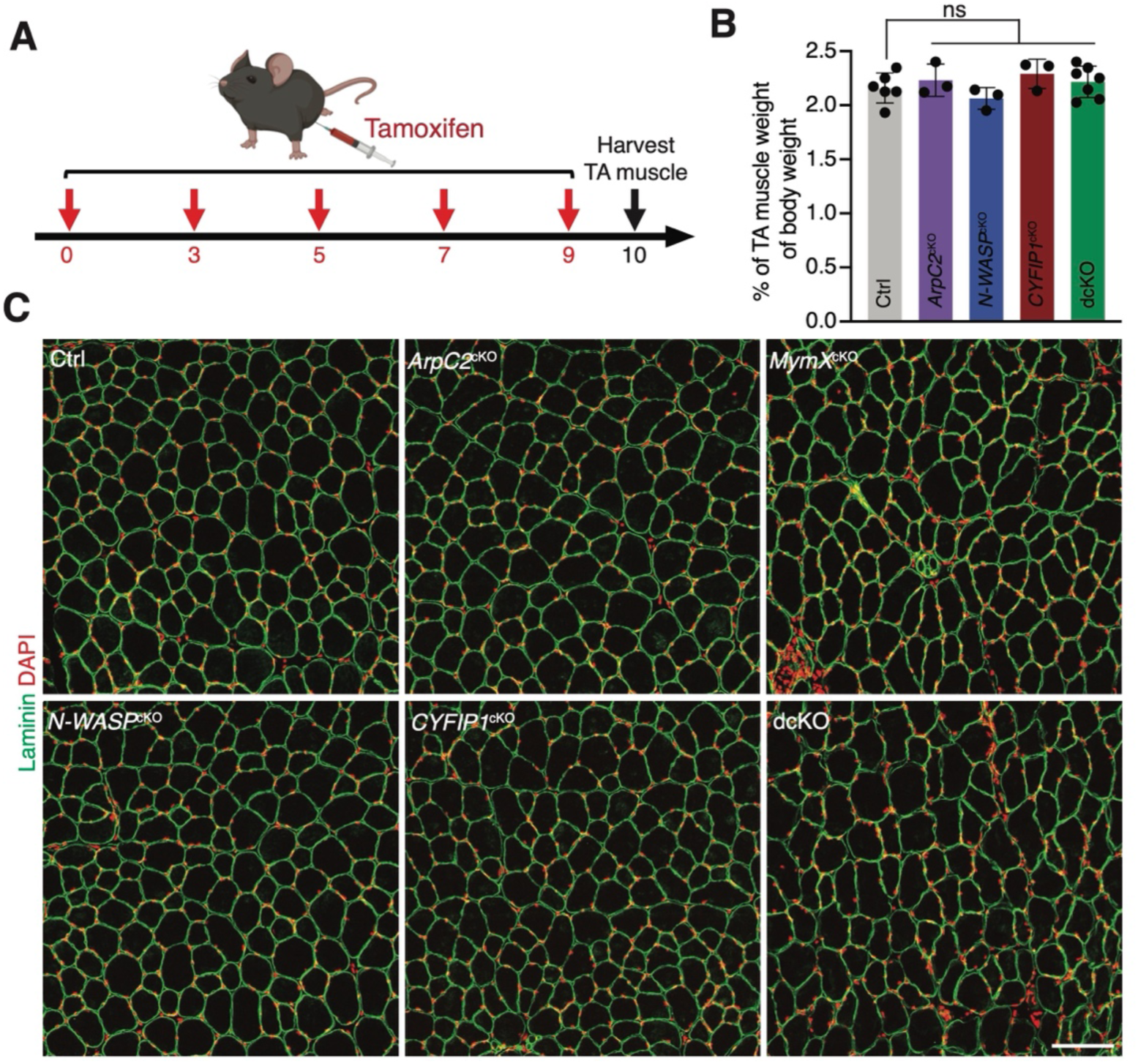
The TA muscle weight and size are not affected by satellite cell-specific deletion of branched actin polymerization regulators before injury. **(A)** Schematic diagram of tamoxifen treatment and TA muscle harvest. **(B)** Quantification of the TA muscle weight. Mice with the indicated genotypes were treated as described in (A). The whole body and TA muscle weights were measured. *n* ≥ 3 mice of each genotype were examined. Mean ± s.d. values are shown in the bar graph, and significance was determined by two-tailed student’s t-test. ns: not significant. **(C)** The TA myofiber size appeared normal in control and mutants with indicated genotypes. Cross sections of TA muscles were stained with anti-Laminin and DAPI. *n* = 3 mice of each genotype were examined with similar results. Scale bar: 100 μm.

**Supplementary Figure 4.**
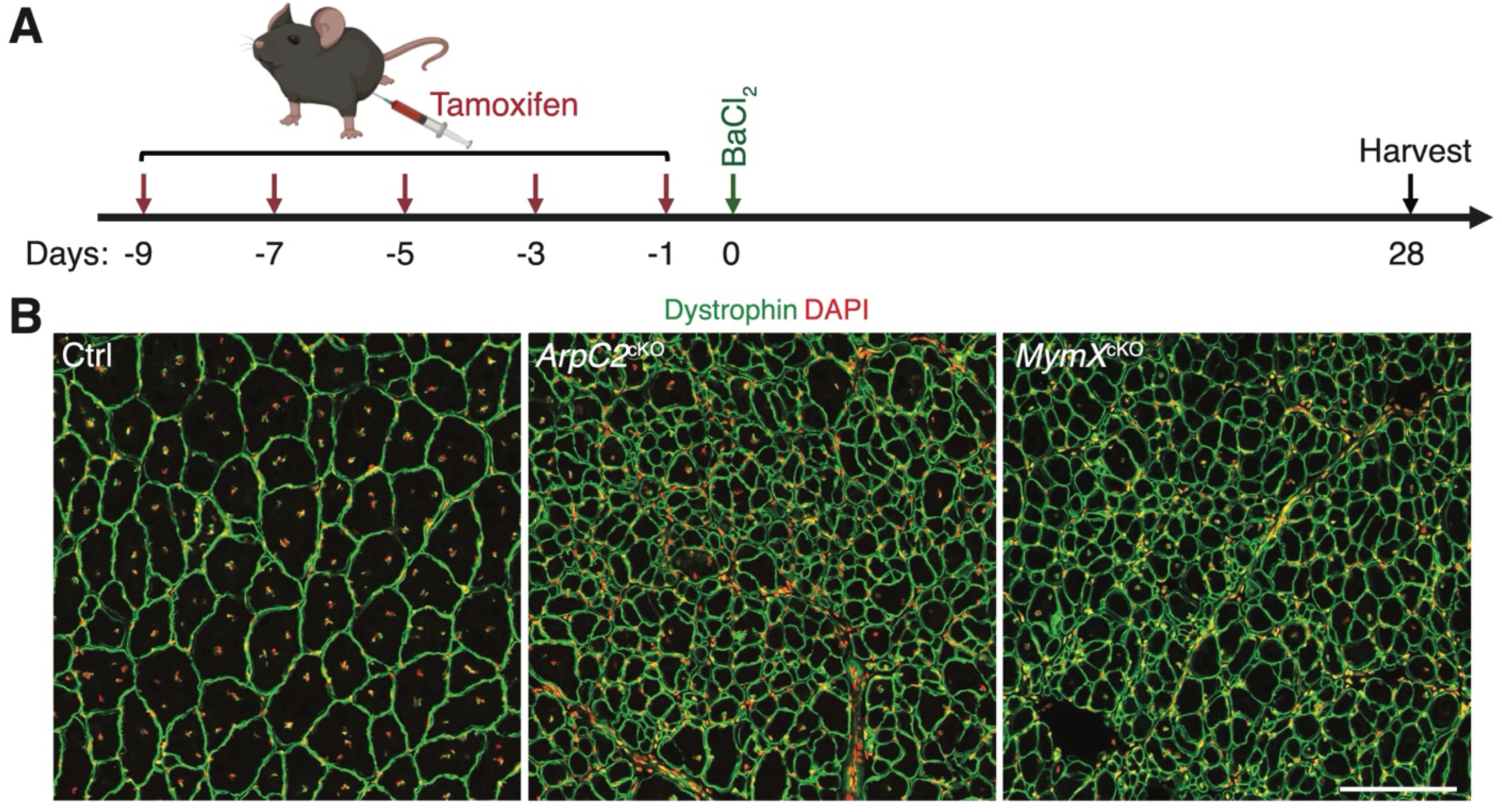
Impaired muscle regeneration in *ArpC2*^cKO^ and *MymX*^cKO^ mice persists to dpi 28. **(A)** Schematic diagram of tamoxifen and BaCI_2_ treatment and subsequent CSA analysis at dpi 28. **(B)** Dystrophin and DAPI staining in TA muscle cross sections at dpi 28 in the control and mutant mice. Note that the myofiber CSA is severely reduced in *ArpC2*^cKO^ and *MymX*^cKO^ mice compared to that of the control mice. Scale bar: 100 μm.

**Supplementary Figure 5.**
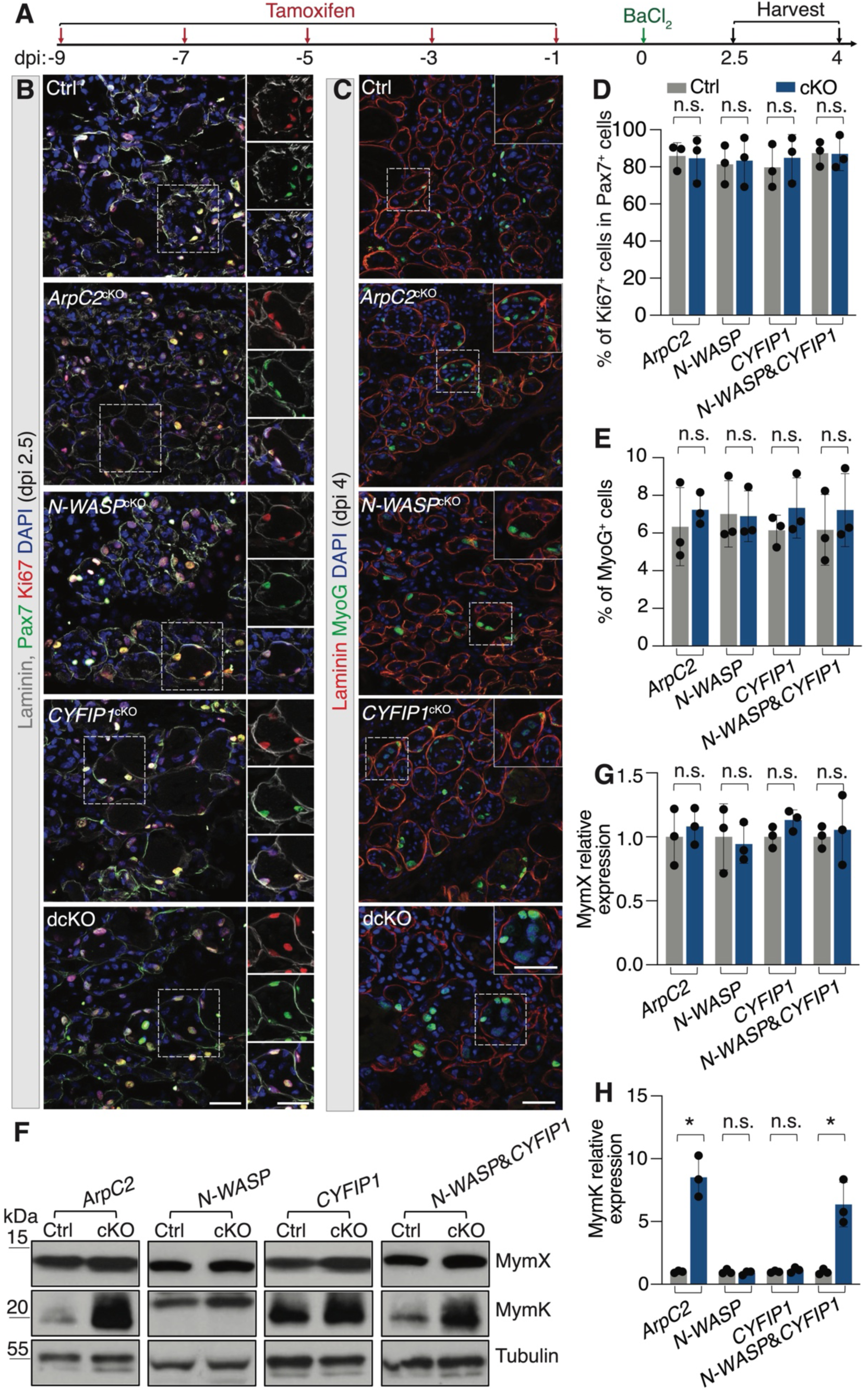
Branched actin polymerization is not required for satellite cell proliferation, differentiation, or fusogenic protein expression. **(A)** Schematic diagram of tamoxifen and BaCI_2_ treatment and subsequent cell proliferation and differentiation analyses at dpi 2.5 and 4, respectively. **(B)** Immunostaining with anti-Laminin, anti-Pax7 and anti-Ki67 of cross sections of TA muscles from control and mutant mice at dpi 2.5. The boxed the areas are shown on the right. Scale bar: 30 μm. **(C)** Immunostaining with anti-Laminin and anti-MyoG of cross sections of TA muscles from control and mutant mice at dpi 4. The boxed areas are shown at the top right corner. Scale bar: 30 μm. **(D)** Quantification of the percentage of proliferating satellite cells (% of Ki67^+^ cells in the Pax7^+^ cells) in TA muscles from control and mutant mice of the indicated genotypes. Mean ± s.d. values are shown in the bar graph, and significance was determined by two-tailed student’s t-test. ns: not significant. **(E)** Quantification of the percentage of MyoG^+^ nuclei in the total cells in TA muscles from control and mutant mice of the indicated genotypes. Mean ± s.d. values are shown in the bar graph, and significance was determined by two-tailed student’s t-test. ns: not significant. **(F)** Western blot analysis for MymX and MymK in TA muscles from control and mutant mice of the indicated genotypes at dpi 4. *n* = 3 mice of each genotype were examined. One sample of each control and mutant genotype were loaded. (**G**, **H**) Quantification of MymX and MymK protein expression in the control and littermate mutant mice as shown in (F). The band intensity of each protein was normalized against β-tubulin. The y axis indicates the expression of MymX or MymK in different mutants relative to the control mice. Mean ± s.d. values are shown in the bar graphs, and significance was determined by two-tailed student’s t-test. *p < 0.05; ns: not significant.

**Supplementary Figure 6.**
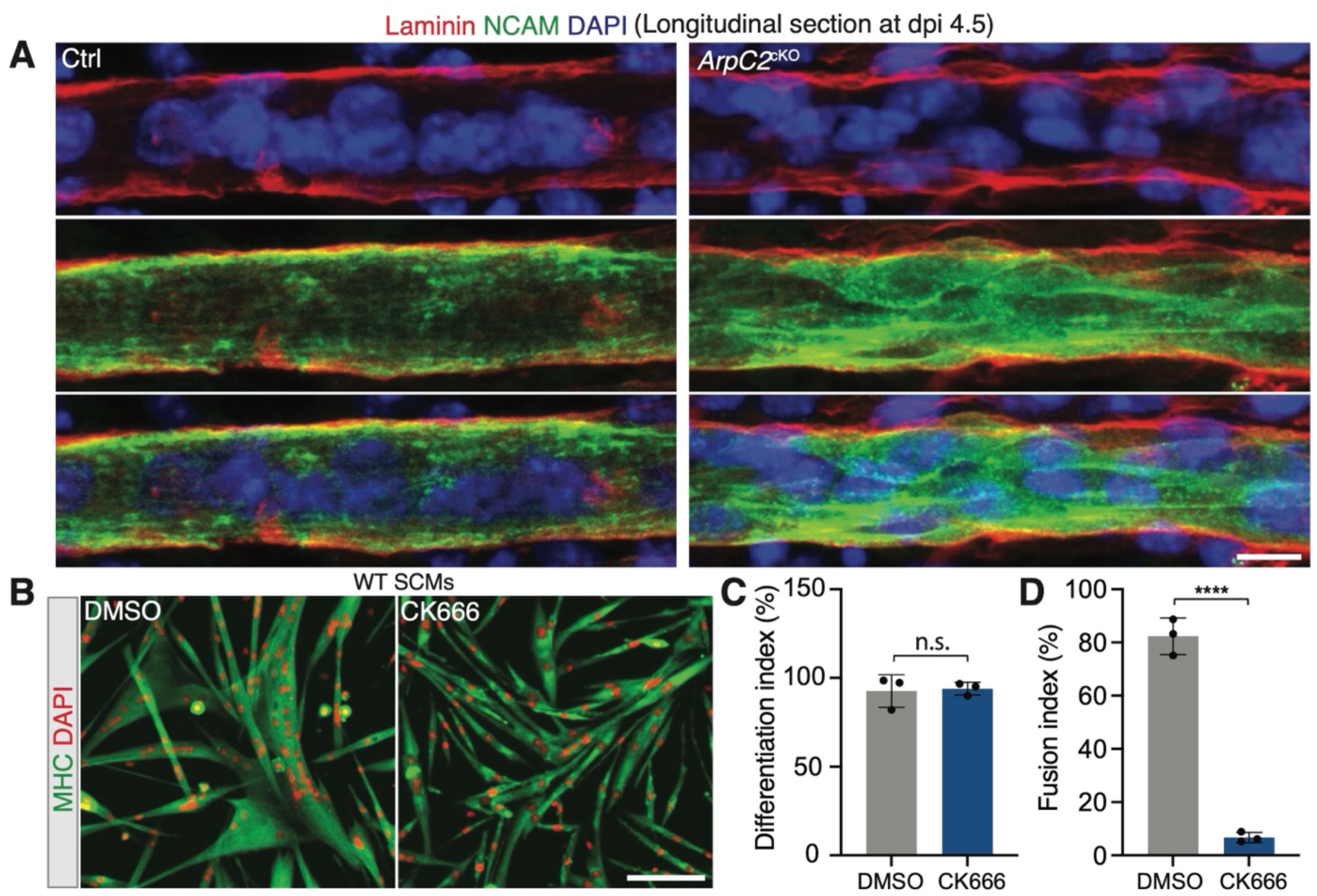
Branched actin polymerization is required for SCM fusion. **(A)** Immunostaining with anti-Laminin and anti-NCAM of longitudinal sections of TA muscles from control and *ArpC2*^cKO^ mice at dpi 4.5. Scale bar: 10 μm. **(B)** Pharmacological inhibition of the Arp2/3 complex during differentiation blocked SCM fusion. The wild-type SCMs were plated at 60% confluence in GM. After a day. The cells were then incubated in DM supplemented with DMSO as a control (0.1%) or the Arp2/3 complex inhibitor CK666 (100 μM) for 48 hours, followed by anti-MHC and DAPI staining. Note the robust fusion of control SCMs *vs*. the severe fusion defects in SCMs treated with CK666. Scale bar: 100 μm. **(C** and **D)** Quantification of the differentiation and fusion indexes of the cells shown in (B). *n* = 3 independent experiments were performed. Mean ± s.d. values are shown in the bar graphs, and significance was determined by two-tailed student’s t-test. ****: p < 0.0001; n.s: not significant.

**Supplementary Figure 7.**
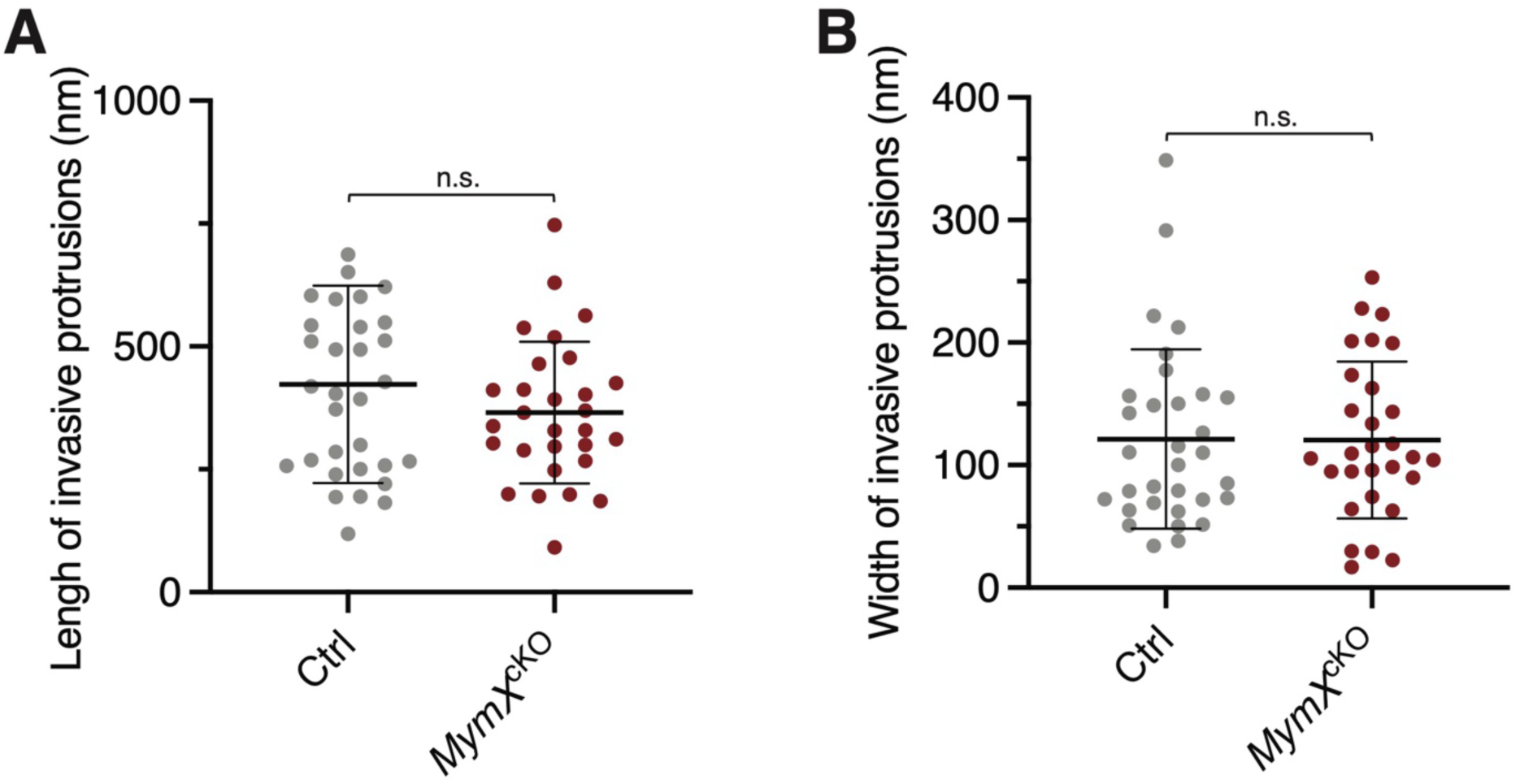
Fusogenic protein MymX is not required for invasive protrusion formation. Quantification of the length (A) and width (B) of the invasive protrusions in control and *MymX*^cKO^ mice at dpi 3.5 imaged by TEM. The width was measured at the midpoints of the invasive protrusions. Mean ± s.d. values are shown in dot plots, and statistical analysis was performed for each parameter in *n* ≥ 29 invasive protrusions in each genotype. Significance was determined by two-tailed student’s t-test. n.s.: not significant.

**Supplementary Figure 8.**
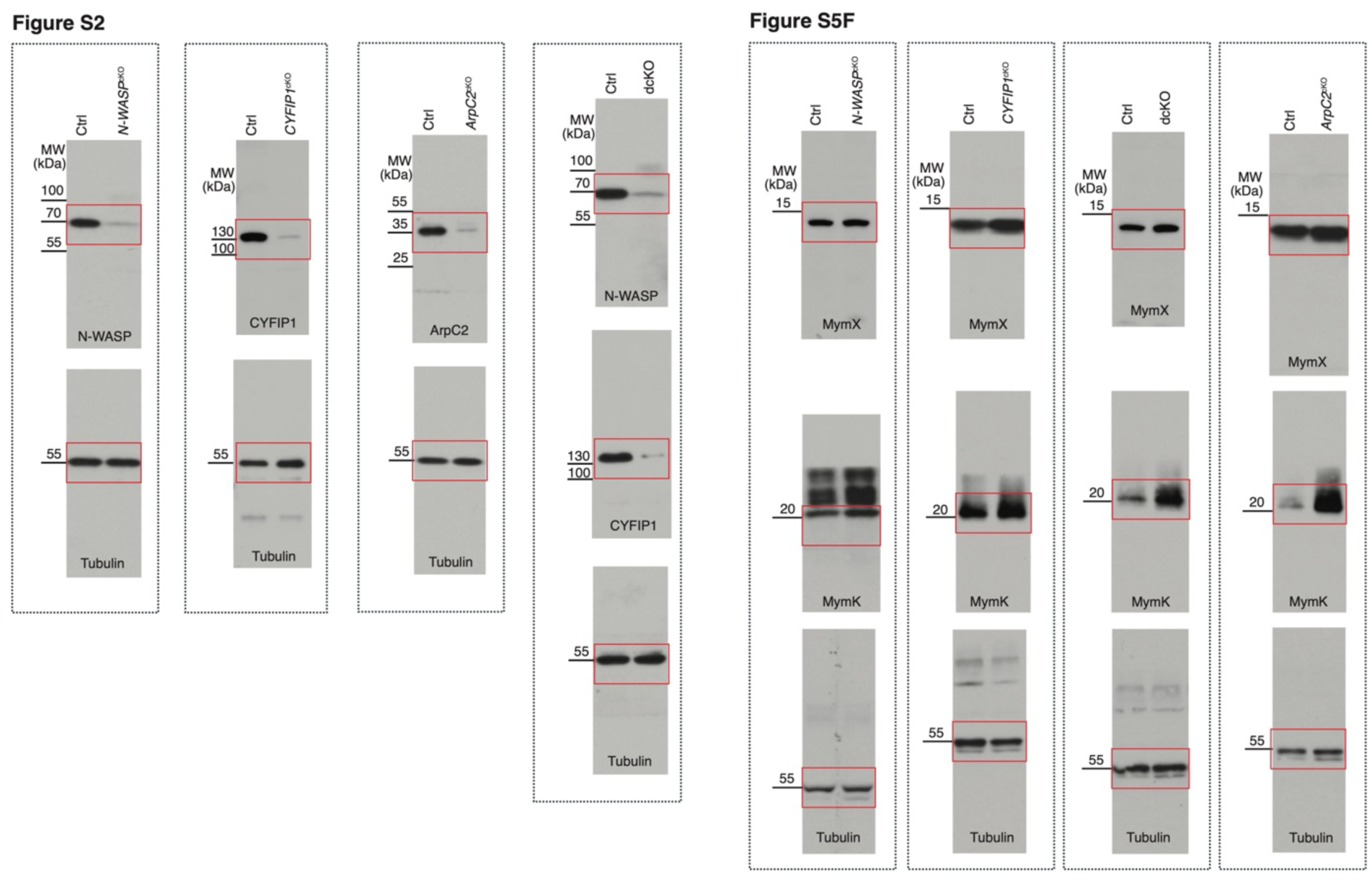
Unprocessed western blot films. The unprocessed western blot films used to generate the Fig. S2 and S5F. The bands shown in Fig. S2 and S5F are highlighted by red boxes.

**Supplemental Video 1. Macrophages extravasate the ghost fibers by traversing the BM– ghost fiber 1**

Representative 3D reconstruction of confocal z-stacks of TA muscle at dpi 3.5. Two examples are shown. The small opening on the BM is indicated by yellow arrowheads, and the transversing macrophage is indicated by magenta arrows.

**Supplemental Video 1. Macrophages extravasate the ghost fibers by traversing the BM– ghost fiber 2**

**Supplemental Video 3. Branched actin polymerization is not required for SCM migration during differentiation.**

Time-lapse imaging of control and *ArpC2* KO SCMs at 24 hours in DM. The SCMs isolated from *ArpC2*^cKO^ mice were maintained in GM without or with 2 μM 4OH-tamoxifen (4OHT) for 10 days. Subsequently, the cells were plated in 70% confluence in GM. After 24 hours, the cells were cultured in DM for 12 hours, followed by live cell imaging. Note that the *ArpC2* KO SCMs were able to migrate normally, although their fusion was significantly impaired. The time interval is five minutes.

**Supplemental Video 4. F-actin and the Arp2/3 complex are enriched in the invasive protrusions at the fusogenic synapse of SCMs**

Time-lapse imaging of a fusion event between two mouse SCMs co-expressing LifeAct-mScar and Arp2-mNG at 24 hours in DM. Note that F-actin and Arp2 were enriched in the finger-like invasive protrusions at the fusogenic synapse (arrows) and dissolved immediately after cell membrane fusion. The time interval is two minutes. Single focal plane is shown.

## Acknowledgements

We thank Dr. Eric Olson for generously providing the *MymX*^fl/fl^ mice, UT Southwestern Animal Resource Center for assistance with mouse colony maintenance, and UT Southwestern Quantitative Light Microscopy Core Facility for assistance with 3D reconstruction of confocal images. This work was supported by an NIH grant (R35GM136316) to E.H.C. Y.L. was supported by an American Heart Association Career Development Award (25CDA1451113).

## Author contributions

Y.L. and E.H.C. designed the project. Y.L., T.W., P.P., and C.Z. performed experiments. Y.L. and E.H.C. collaborated with C.W.H., S.B.S. and R.L. on the mouse genetic experiments. Y.L., and E.H.C. analyzed the data, made the figures, and wrote the manuscript. All authors commented on the manuscript.

## Data availability

The data supporting the findings of this study are available within the article and its supplementary files.

